# The kinesin motor Kif9 regulates centriolar satellite positioning and mitotic progression

**DOI:** 10.1101/2024.04.03.587821

**Authors:** Juan Jesus Vicente, Michael Wagenbach, Justin Decarreau, Alex Zelter, Michael J. MacCoss, Trisha N. Davis, Linda Wordeman

**Author notes:** Correspondence should be addressed to JJV and LW.

## Abstract

Centrosomes are the principal microtubule-organizing centers of the cell and play an essential role in mitotic spindle function. Centrosome biogenesis is achieved by strict control of protein acquisition and phosphorylation prior to mitosis. Defects in this process promote fragmentation of pericentriolar material culminating in multipolar spindles and chromosome missegregation. Centriolar satellites, membrane-less aggrupations of proteins involved in the trafficking of proteins toward and away from the centrosome, are thought to contribute to centrosome biogenesis. Here we show that the microtubule plus-end directed kinesin motor Kif9 localizes to centriolar satellites and regulates their pericentrosomal localization during interphase. Lack of Kif9 leads to aggregation of satellites closer to the centrosome and increased centrosomal protein degradation that disrupts centrosome maturation and results in chromosome congression and segregation defects during mitosis. Our data reveal roles for Kif9 and centriolar satellites in the regulation of cellular proteostasis and mitosis.

## INTRODUCTION

The centrosome is a non-membranous organelle that acts as microtubule-organizing center (MTOC) of the mammalian cell, plays an essential role during the formation of the mitotic spindle in mitosis, it serves as the basal body during cilia formation, and it acts as a hub for protein degradation in the cell ^1–3^. The centrosome is composed of two perpendicular centrioles (cylindrical structure formed by microtubules) surrounded by a matrix of proteins known as the pericentriolar material (PCM) ^1,4,5^. During late G2/early M centrosomes recruit proteins to the PCM where they become increasingly phosphorylated: a process called centrosome maturation ^6^. This process increases the centrosome diameter from ∼500 nm to several micrometers and improves the structural integrity of the centrosome, allowing it to resist the action of spindle forces produced during mitosis ^7,8^. Centriolar satellites (CS) are small (70-100 nm) non-membranous electron-dense granules that accumulate around the centrosomes of animal cells during interphase and regulate protein trafficking to and from this organelle ^9–12^. They disappear during mitosis and reappear upon the return to interphase ^13^. In recent years it has also been suggested that CS regulate cellular proteostasis and protein degradation at the centrosome by spatially regulating access of key proteolysis regulators to the centrosome ^14^. CS settle at a distance from the centrosome in a microtubule (MT) dependent manner ^15^, and while it is known that dynein regulates the movement of CS towards the centrosome, the identity of the molecular motor directing CS towards the plus end of the MT is not known ^16^.

Kif9, a member of the Kinesin-9 family, was originally identified as Klp1 in the *Chlamydomonas* flagellum in 1994 ^17^. In mammals, Kif9 expression is dominant in testis, with lower expression in kidney, spleen and lung ^18^. Its activity has been linked to mRNA transport ^19^, macrophage podosome homeostasis ^20^, and mitosis ^21^. It has also appeared in screens related to breast cancer ^22^, glioblastoma ^23^, mouse inner ear development ^24^, nuclear movement during skeletal muscle differentiation ^25^, and sperm flagellum formation and maintenance ^26,27^. However, Kif9’s fundamental cellular function remains poorly understood. In this study we show that Kif9, a plus-end directed motor, localizes to CS and is responsible for setting their pericentrosomal positioning during interphase. Loss of Kif9, in turn, promotes centrosomal protein degradation resulting in profound mitotic defects.

## RESULTS

### Kif9 is a plus-end directed motor that localizes to centriolar satellites

Human Kif9 has 790 amino acids with a motor domain positioned at the N-terminus of the protein, followed by a dimerization domain in the central stalk region and a C-terminal tail ^28^. This motor domain localization at the N-terminal would suggest plus-end directionality on the microtubule. Using purified eGFP-Kif9 and total internal reflection fluorescence (TIRF) microscopy we have confirmed that Kif9 is a plus-end directed motor with an average speed of 10.40 ± 0.21 nm/sec (mean ± s.e.m.) (Fig. 1A and 1B). Similar *in vitro* speeds have been reported recently ^29^. Kif9 localizes to small granules around the centrosome in interphase HeLa cells (Fig. 1C), HCT116 cells (Supplementary Fig. 1A) and also in CRISPR knock-in HCT116 cell lines where Kif9 is fused to either eGFP or Neon-Green, wherein it is expressed under its endogenous promoter (Supplementary Fig. 1B and C respectively). To ascertain the interacting partners of Kif9 we performed mass spectrometry of protein samples pulled down with an anti-GFP antibody from HCT116 cells expressing eGFP-Kif9 versus control HCT116 cells. Table I shows that the centriolar satellites marker PCM1 is the principal Kif9 interactor in interphase cells (Table I). PCM1 is a 228 kDa protein that acts as a structural platform for CS assembly, and the first protein identified to be associated with CS ^16,30^. Accordingly, it is generally accepted that a protein localizes to CS if it co-localizes with PCM1. To confirm the interaction between Kif9 and PCM1 we first used fluorescence microscopy to show that a majority of eGFP-Kif9 spots co-localized with PCM1 positive spots in fixed HeLa cells (Fig. 1D). Using the software CellProfiler ^31^ we quantified the fluorescence overlap between spots found in the Kif9 and PCM1 channels. Manders’ overlap coefficient showed that 77% of the Kif9 spots overlap with PCM1 spots, whereas 43% of PCM1 spots overlap with Kif9 (Fig. 1E). Published proteomic data has shown that the protein composition of individual CS is variable, making it possible that some PCM1 granules do not interact with Kif9 ^32,33^. Interestingly, our mass spectrometry analysis did not show enrichment of dynein in the eGFP tagged Kif9 pulldowns over the control samples, suggesting that CS may not interact with both motors at the same time (Table I). Super resolution Structural Illumination Microscopy (SIM) also confirmed the co-localization between Kif9 and PCM1 (Supplementary Fig. 1D). Immunoprecipitation was used to further verify the direct interaction of Kif9 and PCM1. HeLa cells were transfected with either eGFP-empty or eGFP-Kif9 plasmids followed by immunoprecipitation with an anti-GFP antibody and western-blot to detect PCM1. Lysate from cells expressing eGFP-Kif9 co-immunoprecipitated with a PCM1 positive band (228 kDa predicted molecular weight), confirming the interaction between these proteins (Fig. 1F). Finally, we used a knock-sideways approach where Kif9 fused to FKBP (GFP-FK506–binding protein ^34^) associates with regions of the cell where we have anchored FRB (FKBP12 Rapamycin binding domain of mTOR) in the presence of rapamycin. In our experiments, FRB was fused to blue fluorescent protein (BFP) and to the membrane-bound domain SH4. PCM1 remain pericentrosomal in the absence of Rapamycin or without SH4-FRB-BFP. However, in cells expressing both plasmids (eGFP-FKBP-Kif9 + SH4-FRB-BFP), Kif9 and PCM1 granules move away from the centrosomal area and relocalize to regions close to the cell membrane in the presence of Rapamycin (Fig. 2A). These three approaches indicate that Kif9 interacts with PCM1, the major protein and marker for CS.

**Figure 1.**
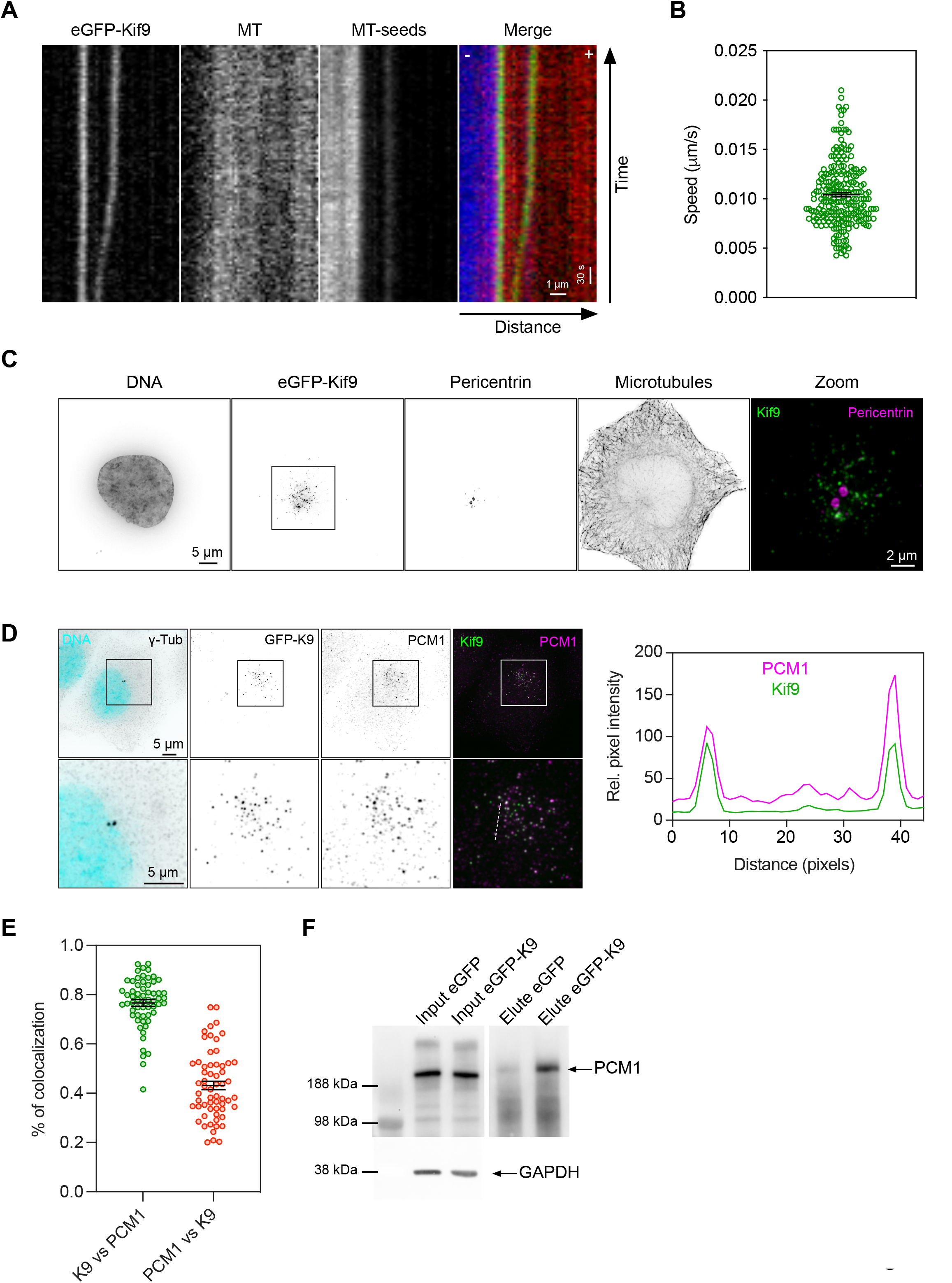
Kif9 is a microtubule (MT) plus-end directed motor that localizes to centriolar satellites (CS). **(A)** Kymograph showing movement of eGFP-Kif9 on dynamic MTs towards the MT plus-end (the right side of the panel). **(B)** Quantification of eGFP-Kif9 events speed on live MTs. The calculated mean speed for Kif9 was 10.40 ± 0.21 nm/sec (mean ± s.e.m.). Data is from n = 233 moving spots pooled from 17 movies obtained in two independent experiments. **(C)** eGFP-Kif9 localizes to spot-like structures around the centrosome. HeLa cells were transfected with a plasmid for eGFP-Kif9 expression, fixed and stained with DAPI for DNA, and antibodies against Pericentrin (for centrosomes) and tubulin (for microtubules). Scale bar, 5 µm for the regular panels and 2 µm for the zoom panel. **(D)** Kif9 co-localizes with the CS marker PCM1. HeLa cells were transfected with a eGFP-Kif9 plasmid, fixed and stained with DAPI for DNA, and antibodies against PCM1 (for CS) and ɣ-tubulin (for centrosomes). Bottom panels are magnifications of the square in top panels. Scale bar, 5 µm for all panels. Graph on the right is a plot profile of the line in the magnified merge channel (bottom right panel). **(E)** Kif9-PCM1 co-localization quantification using Manders’ overlap coefficient. Mean value for “K9 vs PCM1” is 0.766 ± 0.0132 and for “PCM1 vs K9” 0.431± 0.018 (mean ± s.e.m.). Data is from n = 60 cells from three independent experiments. **(F)** Co-immunoprecipitation Kif9-PCM1. Immunoprecipitation of eGFP-Kif9 shows an interaction with PCM1. HeLa cells were transfected either with eGFP-Kif9 or the eGFP empty plasmid. Immunoprecipitation was performed with an anti-GFP antibody followed by immunoblotting with an anti-PCM1 antibody.

**Figure 2.**
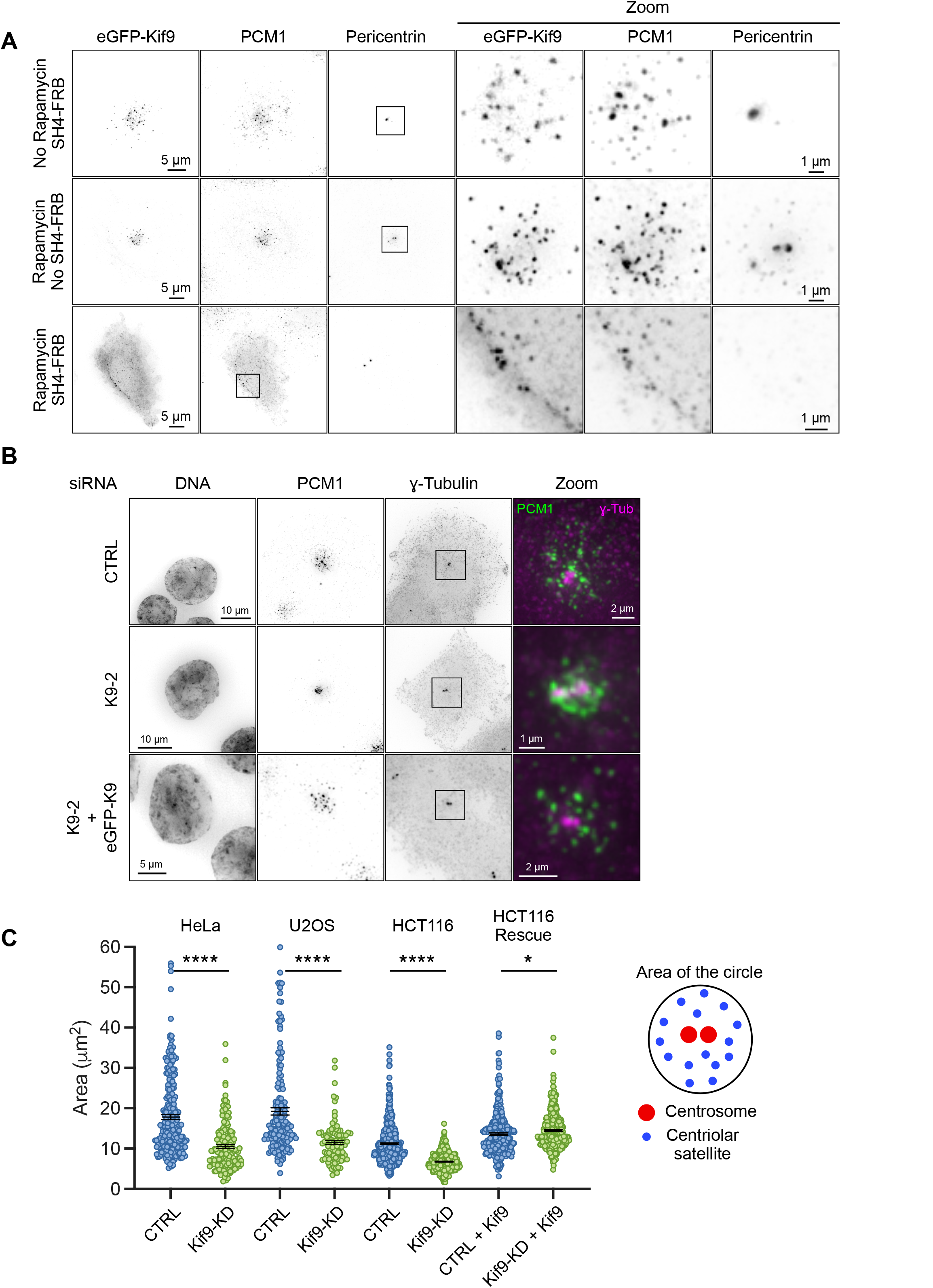
Lack of Kif9 or changes in its location affects CS positioning. **(A)** HeLa cells were transfected with either eGFP-FKBP-Kif9 (relocalizable form of Kif9) or eGFP-FKBP-Kif9 plus SH4-FRB-BFP. The co-transfection with SH4-FRB-BFP allows the relocalization of Kif9 from internal structured to the cell membrane after treatment with Rapamycin. The top panels show a cell transfected with both construction but without Rapamycin. In this situation CS keep their localization around the centrosome. The middle panels show a cell transfected with eGFP-FKBP-Kif9, treated with 1 µM Rapamycin O/N but without FRB-BFP at the cell membrane. CS are still around the centrosome after treatment. The bottom panels show a cell transfected with eGFP-FKBP-Kif9, FRB-BFP at the cell membrane and treated with 1 µM Rapamycin O/N. In this case we can see how the CS (labeled with PCM1) moved away from the centrosome and co-localize with the Kif9 spots (bottom inset magnifications). Scale bar, 5 µm for the regular panels and 1 µm for the zoom panels. Images are fixed cells stained with DAPI for DNA, and antibodies against PCM1 (for CS) and Pericentrin (for centrosomes). **(B)** HCT116 cells treated with a control siRNA (top panels), Kif9 siRNA (middle panel) or Kif9 siRNA plus co-transfection with an siRNA resistant form of eGFP-Kif9 were fixed and stained with DAPI for DNA, and antibodies against PCM1 (for CS) and ɣ-tubulin (for centrosomes). Lack of Kif9 results in CS closer to the centrosome (middle panels); phenotype recovered by over-expression of Kif9 (bottom panels). Scale bar: top panels, 10 µm for the regular panels and 2 µm for the zoom panel; middle panels, 10 µm for the regular panels and 1 µm for the zoom panel; bottom panels, 5 µm for the regular panels and 2 µm for the zoom panel. **(C)** Quantification of CS distance from the centrosome in different cell lines depleted for Kif9 (HeLa, U2OS and HCT116). Cells were fixed and stained with DAPI for DNA, and antibodies against PCM1 (for CS) and ɣ-tubulin (for centrosomes). We quantified the positioning of CS drawing a circle around them and measuring the area using Fiji. For the rescue experiment we co-transfected the siRNAs with a plasmid expressing a siRNA resistant form of eGFP-Kif9. Data is from n (number of cells) = 235 (HeLa CTRL), 164 (HeLa K9KD), 158 (U2OS CTRL), 98 (U2OS K9KD), 483 (HCT116 CTRL), 423 (HCT116 K9KD), 316 (HCT116 CTRL+Kif9), and 424 (HCT116 K9KD+Kif9) from 3 independent experiments (HeLa and HCT116) and 2 independent experiments (U2OS). Bars are mean ± s.e.m. We used an unpaired t test as statistical analysis and the GraphPad Prism nomenclature where p values are categorized as follow 0.1234 (ns), 0.0332 (*), 0.0021 (**), 0.0002 (***), <0.0001 (****).

**Table I.**
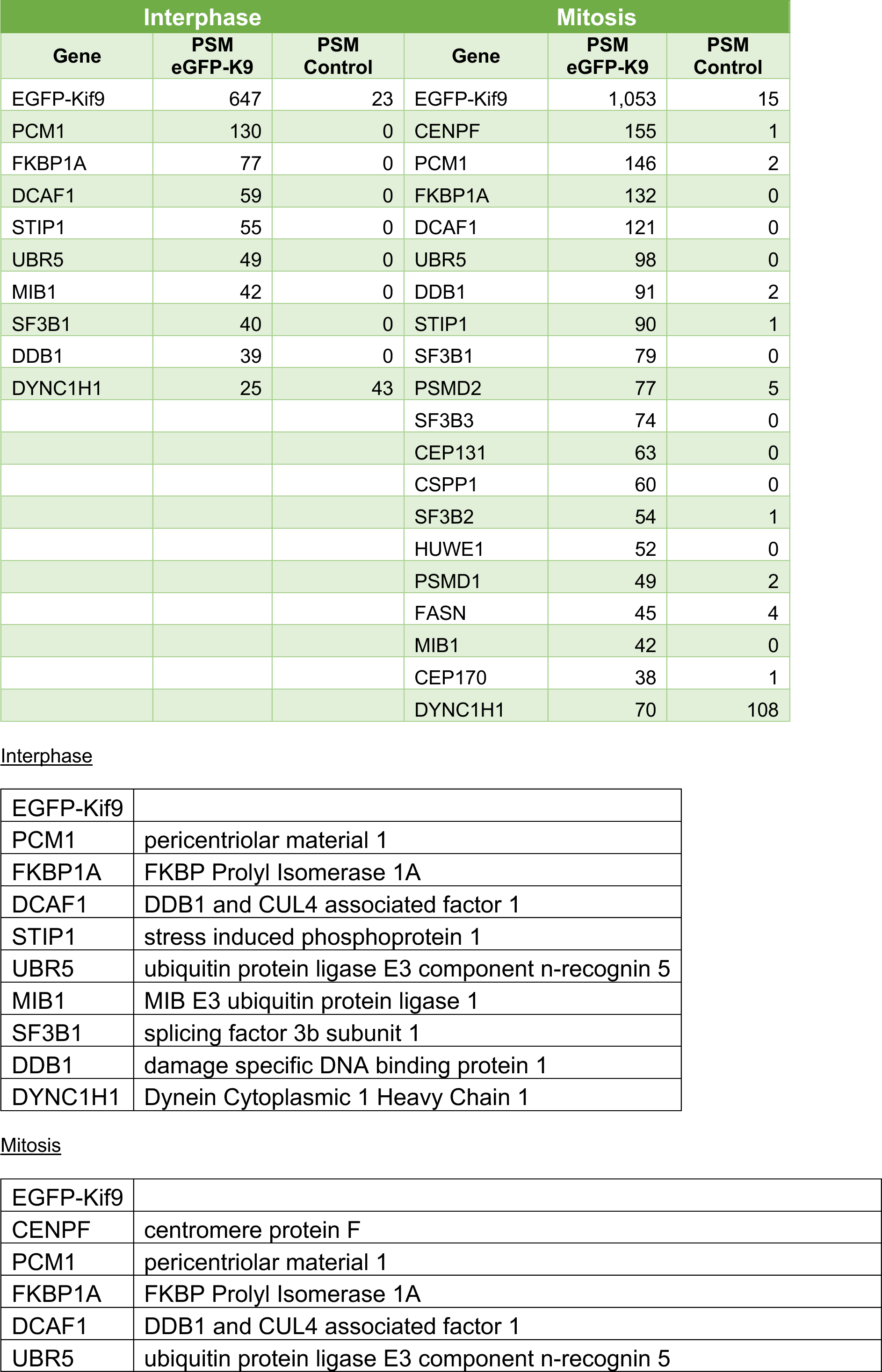
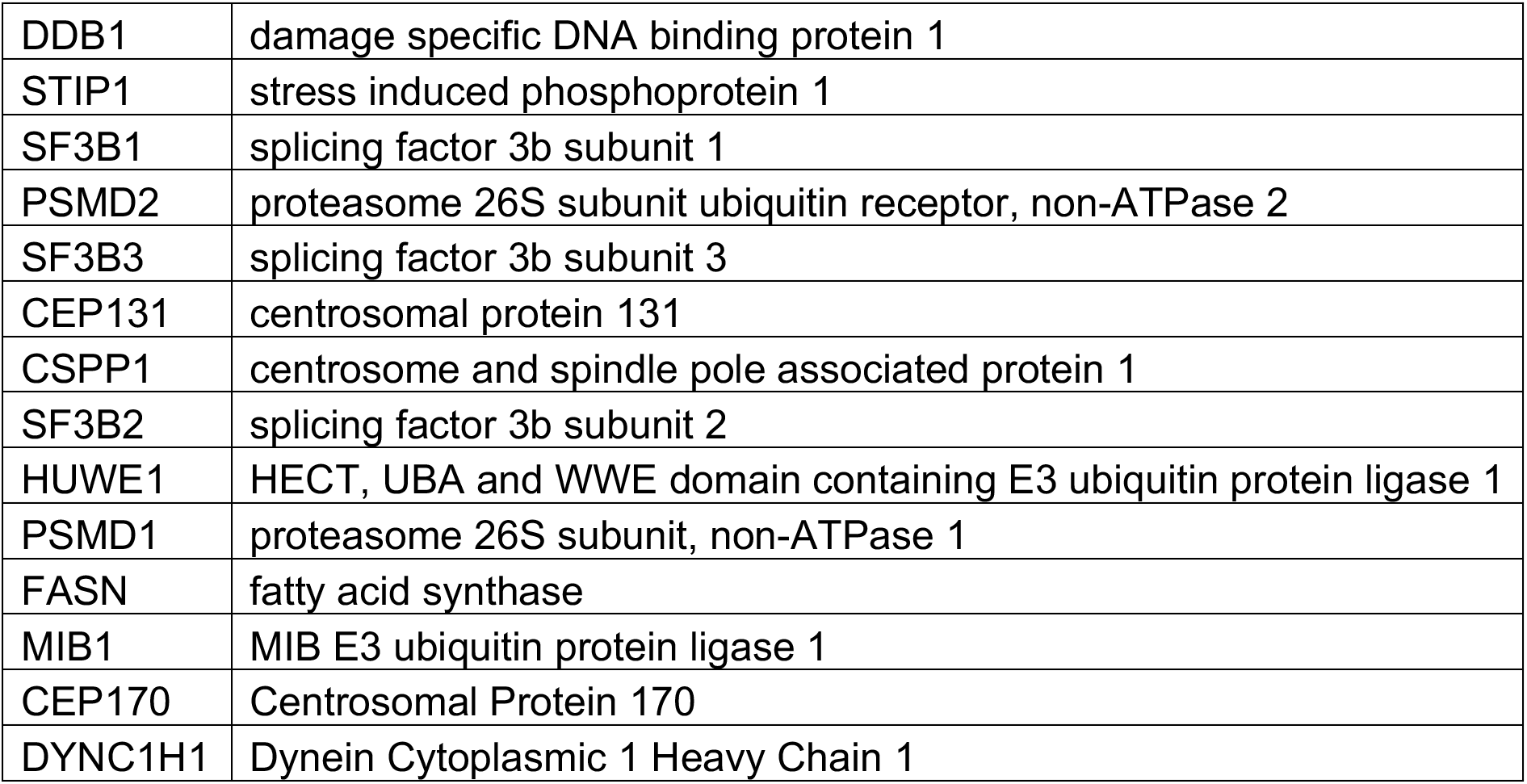
Selected hits from mass spectrometry analysis from HCT116-eGFP-Kif9 or HCT116 cells. The columns show the PSM (peptide spectrum matches) values for control (HCT116 cells) and Kif9 (eGFP-Kif9) cells. Cells treated or not with colcemid provided information for mitotic and interphase cells respectively. Proteins were filtered doing a ratio dividing the values of PSMs-CTRL by the values of PSMs-eGFP-Kif9 and removing all those hits with values over 0.1 (i.e. we kept those hits with at least a 10:1 ratio eGFP-K9:CTRL). Afterwards we removed all those hits with PSMs values for eGFP-Kif9 below 35. Where there were multiple protein homologues that shared all or the majority of identified peptides, only the homologue with the most identified peptides was shown for simplicity. Dynein, a previously known CS component, is included in the table to show the lack of enrichment in eGFP tagged Kif9 pulldowns over the control samples.

**Supplementary Figure 5**
| Interphase |  |  | Mitosis |  |  |
| --- | --- | --- | --- | --- | --- |
| Gene | PSM<br>eGFP-K9 | PSM<br>Control | Gene | PSM<br>eGFP-K9 | PSM<br>Control |
| EGFP-Kif9 | 647 | 23 | EGFP-Kif9 | 1,053 | 15 |
| PCM1 | 130 | 0 | CENPF | 155 | 1 |
| FKBP1A | 77 | 0 | PCM1 | 146 | 2 |
| DCAF1 | 59 | 0 | FKBP1A | 132 | 0 |
| STIP1 | 55 | 0 | DCAF1 | 121 | 0 |
| UBR5 | 49 | 0 | UBR5 | 98 | 0 |
| MIB1 | 42 | 0 | DDB1 | 91 | 2 |
| SF3B1 | 40 | 0 | STIP1 | 90 | 1 |
| DDB1 | 39 | 0 | SF3B1 | 79 | 0 |
| DYNC1H1 | 25 | 43 | PSMD2 | 77 | 5 |
|  |  |  | SF3B3 | 74 | 0 |
|  |  |  | CEP131 | 63 | 0 |
|  |  |  | CSPP1 | 60 | 0 |
|  |  |  | SF3B2 | 54 | 1 |
|  |  |  | HUWE1 | 52 | 0 |
|  |  |  | PSMD1 | 49 | 2 |
|  |  |  | FASN | 45 | 4 |
|  |  |  | MIB1 | 42 | 0 |
|  |  |  | CEP170 | 38 | 1 |
|  |  |  | DYNC1H1 | 70 | 108 |

Interphase
|  |  |
| --- | --- |
| EGFP-Kif9 |  |
| PCM1 | pericentriolar material 1 |
| FKBP1A | FKBP Prolyl Isomerase 1A |
| DCAF1 | DDB1 and CUL4 associated factor 1 |
| STIP1 | stress induced phosphoprotein 1 |
| UBR5 | ubiquitin protein ligase E3 component n-recognin 5 |
| MIB1 | MIB E3 ubiquitin protein ligase 1 |
| SF3B1 | splicing factor 3b subunit 1 |
| DDB1 | damage specific DNA binding protein 1 |
| DYNC1H1 | Dynein Cytoplasmic 1 Heavy Chain 1 |

Mitosis
|  |  |
| --- | --- |
| EGFP-Kif9 |  |
| CENPF | centromere protein F |
| PCM1 | pericentriolar material 1 |
| FKBP1A | FKBP Prolyl Isomerase 1A |
| DCAF1 | DDB1 and CUL4 associated factor 1 |
| UBR5 | ubiquitin protein ligase E3 component n-recognin 5 |
| DDB1 | damage specific DNA binding protein 1 |
| STIP1 | stress induced phosphoprotein 1 |
| SF3B1 | splicing factor 3b subunit 1 |
| PSMD2 | proteasome 26S subunit ubiquitin receptor, non-ATPase 2 |
| SF3B3 | splicing factor 3b subunit 3 |
| CEP131 | centrosomal protein 131 |
| CSPP1 | centrosome and spindle pole associated protein 1 |
| SF3B2 | splicing factor 3b subunit 2 |
| HUWE1 | HECT, UBA and WWE domain containing E3 ubiquitin protein ligase 1 |
| PSMD1 | proteasome 26S subunit, non-ATPase 1 |
| FASN | fatty acid synthase |
| MIB1 | MIB E3 ubiquitin protein ligase 1 |
| CEP170 | Centrosomal Protein 170 |
| DYNC1H1 | Dynein Cytoplasmic 1 Heavy Chain 1 |

### Loss of Kif9 results in centriolar satellites positioned closer to the centrosome

To investigate the function of Kif9 and its role in CS homeostasis we performed RNA interference (RNAi) experiments. Treatment of HeLa cells with three different siRNAs against Kif9 reduces Kif9 mRNA levels as measured by Q-RT-PCR (Supplementary Fig. 2A-D). SiRNAs number 2 and 4 caused the most pronounced reduction on mRNA levels, and therefore, we proceeded with these two siRNAs for the study. HCT116 cells knocked-down for Kif9 presented CS occupying a smaller area around the centrosome (Fig. 2B-C and Supplementary Fig. 2E). The same phenotype was evident in HeLa and U2OS cells (Fig. 2C). Moreover, HCT116 cells knocked-down for Kif9 and transfected with an eGFP-Kif9 plasmid resistant to siRNAs exhibited rescue of the phenotype and a return of CS to their normal position (Fig. 2B-C). Altogether, these data suggest that Kif9 is a PCM1 interactor and has a role regulating the positioning of CS relative to the centrosome.

### A plethora of mitotic defects appear in cells lacking Kif9

Cells depleted of Kif9 also show defects in chromosome congression, segregation, and exhibit fragmentation of the pericentriolar matrix (PCM) (Fig. 3A-C). While control cells present PCM fragmentation in just ∼5% of the mitotic cells, cells depleted for Kif9 showed fragmentation in ∼88% of the cells (Fig. 3B). Also, fifty-five percent of Kif9-KD anaphase cells showed lagging chromosomes compared to the 3% seen in control cells (Fig. 3C-D, Supplementary Fig. 3A). These phenotypes are reflected in the spindle profile, where Kif9-KD cells showed an increase in the percentage of cells in prometaphase and a decrease in the cells in metaphase (Fig. 3D and Supplementary Fig. 3B). We also see doubling of the mitotic index from ∼3.5% in control cells to ∼7.7% in Kif9-KD cells (Supplementary Fig. 3C). Increases in the mitotic index and percentage of cells in prometaphase, along with congression defects suggests an increase in mitotic timing. To confirm this, we used a stable HeLa cell line expressing the histone H2B fused to GFP (HeLa-H2B-GFP) to follow cells through mitosis and measure the timing from nuclear envelope breakdown (NEB) to anaphase onset (AO). Control cells were able to congress the chromosomes properly and had a mean time from NEB to AO of ∼52 min with most of the cells completing mitosis between 25 and 75 min (Fig. 3E-F and Supplementary figure 3D). Meanwhile, Kif9-KD cells showed congression problems, missegregation of chromosomes during anaphase and exhibited micronuclei after mitosis (Fig. 3E). Kif9 depleted cells exited mitosis with an average time from NEB to AO of ∼108 minutes, with most cells completing mitosis with a timing between 50 and 125 minutes (Fig. 3F and Supplementary Fig. 3D). We saw similar phenotypes for the siRNA#4 (Supplementary Fig. 3D-F) and in HCT116 cells (Supplementary Fig. 3H). The congression defects could not be attributable to loss of the chromosome alignment motor Cenp-E ^35^ as it properly localizes to kinetochores in Kif9 depleted cells (Supplementary Fig. 3G). These data suggest that the lack of Kif9, and accordant positioning of CS closer to the centrosome, adversely affects centrosome structure/homeostasis to trigger mitotic defects.

**Figure 3.**
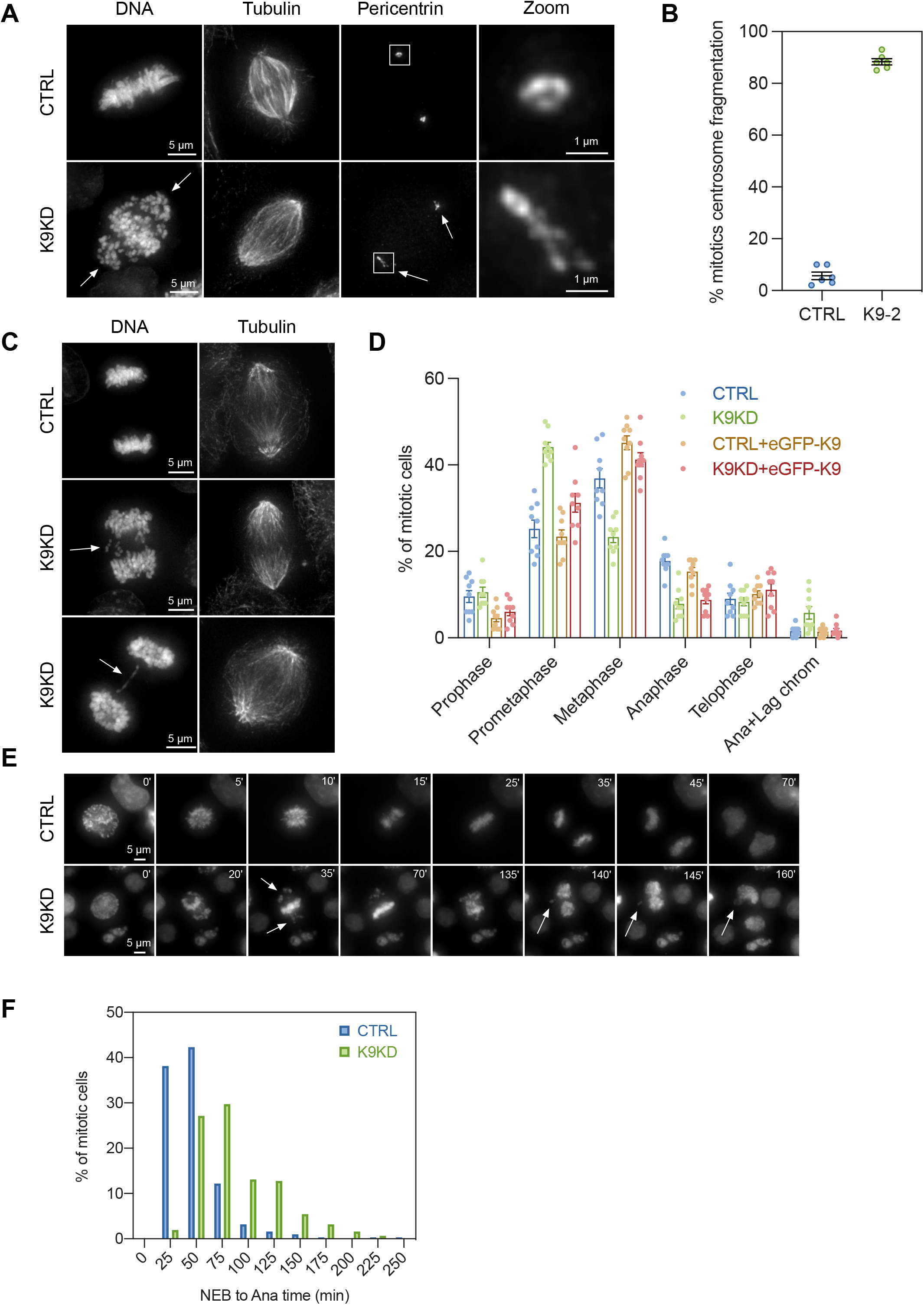
Kif9 depletion induces PCM fragmentation, and chromosome congression and segregation problems during mitosis. **(A)** HeLa cells were transfected with either control or Kif9 siRNAs, fixed and stained with DAPI for DNA, and antibodies against tubulin (for MTs) and Pericentrin (for centrosomes). Kif9 depleted cells showed congression problems (arrows in DNA channel) and fragmentation of the PCM (arrows in Pericentrin channel). The zoom panels are magnifications of the white squares in the Pericentrin channel. Scale bar: 5 µm for the regular panels and 1 µm for the zoom panels. **(B)** Quantification of PCM fragmentation in HeLa mitotic cells transfected with control or Kif9 siRNAs from the figure (a). n = 600 mitotic cells per condition from 3 independent experiments. Bar is expressed as mean ± s.e.m. (5.667 for control and 88.33 for Kif9-KD). **(C)** HeLa cells were transfected with either control or Kif9 siRNAs, fixed and stained with DAPI for DNA, and antibodies against tubulin (for MTs). Kif9 depleted cells showed lagging chromosomes and chromosomes bridges during anaphase (arrows in DNA channel). Scale bar, 5 µm. **(D)** Quantification of the percentage of cells in every mitotic stage in fixed cells (spindle profile). HeLa cells were transfected with control siRNA, Kif9 siRNA, or co-transfected with siRNAs and a form of eGFP-Kif9 resistant to siRNAs. Cells were then fixed and stained with DAPI for DNA, and antibodies against tubulin (for MTs) and Pericentrin (for centrosomes). Kif9 depleted cells showed an increase in the number of prometaphase and a decrease of metaphases, phenotype recovered by the expression of eGFP-Kif9 siRNA resistant. Cells lacking Kif9 also showed increased levels of anaphases with lagging chromosomes, also recovered by the expression of eGFP-Kif9. n = 900 mitotic cells per condition from 3 independent experiments. Bars are mean ± s.e.m. **(E)** Mitotic timing from nuclear envelope breakdown (NEB) to anaphase onset (AO). Stable HeLa cells expressing H2B-GFP were transfected with either control or Kif9 siRNAs, and recorded for 24 hours with 5 minutes steps. Representative images of the time lapse showing that Kif9 depleted cells have increased mitotic timing with chromosome congression and segregation problems, and the formation of small independent nuclei after nuclear formation (arrows in Kif9-KD panels). Scale bar, 5 µm. **(F)** Histogram of the quantification of mitotic timing movies from (e). Kif9 depleted cells show increased mitotic timing. Data from n = 312 control and 313 Kif9-KD cells from 3 independent experiments.

### Mispositioning of centriolar satellites affects centrosomal proteins levels

PCM fragmentation during mitosis occurs in immature centrosomes that have not acquired the proper set of proteins during late G2/early M. These include the kinases PLK1 (Polo-Like Kinase 1) and AURKA (Aurora Kinase A) ^36^. Plk1 is essential for the acquisition and phosphorylation of PCM proteins and for PCM expansion that provide strength and ductility to the PCM ^37–42^. This resistance to deformation is essential for a structurally stable centrosome that is able to resist MT-dependent pulling forces from the spindle. Loss of Plk1 activity during mitosis leads to rapid depletion of PCM components, followed by deformation and loss of structural integrity of the PCM ^43,44^. Aurora Kinase A phosphorylates Plk1 triggering its activation and promoting its recruitment to the centrosomes ^45^. AURKA also promotes the recruitment of MT-nucleating proteins and several other proteins required for PCM expansion and maturation in preparation for mitosis ^46^. Accordingly, we measured PLK1 and AURKA levels at the centrosome in mitotic cells depleted of Kif9. They presented lower levels of both proteins compared to control (Fig. 4A-B). Concomitant with this reduction, we also saw lower levels of P-PLK1 and P-AURKA in Kif9 depleted cells at the centrosomes during mitosis (Supplementary Fig. 4A-B). AURKA is also essential for the recruitment of ɣ-tubulin to the PCM and predictably Kif9-KD cells also exhibited decreased ɣ-tubulin at the centrosome during mitosis (Fig. 5A). Centrosomal protein Cep170 specifically binds to the subdistal appendages of the parent centriole, interacts with PLK1 during mitosis ^47^ and has been implicated in targeting motors to the mitotic spindle ^48^. Interestingly, Cep170 also localizes to CS in Hek293T and Jurkat cells ^32^. Cep170 reduction at centrosome during mitosis also promotes chromatin alignment defects similar to those that we have seen in the Kif9-KD ^49^. Cep170 appears in our mass spectrometry data as a Kif9 interactor in mitosis (Table I). We confirmed this interaction via immunoprecipitation (Supplementary Fig. 4C) and found that depletion of Kif9 in HeLa cells also reduces the levels of Cep170 at the mitotic centrosome (Supplementary Fig. 4D). Thus, our data suggests that when CS are closer to the centrosome, maturation is defective due to decreased levels of key centrosomal players, triggering a set of mitotic defects that arise from PCM fragmentation.

**Figure 4.**
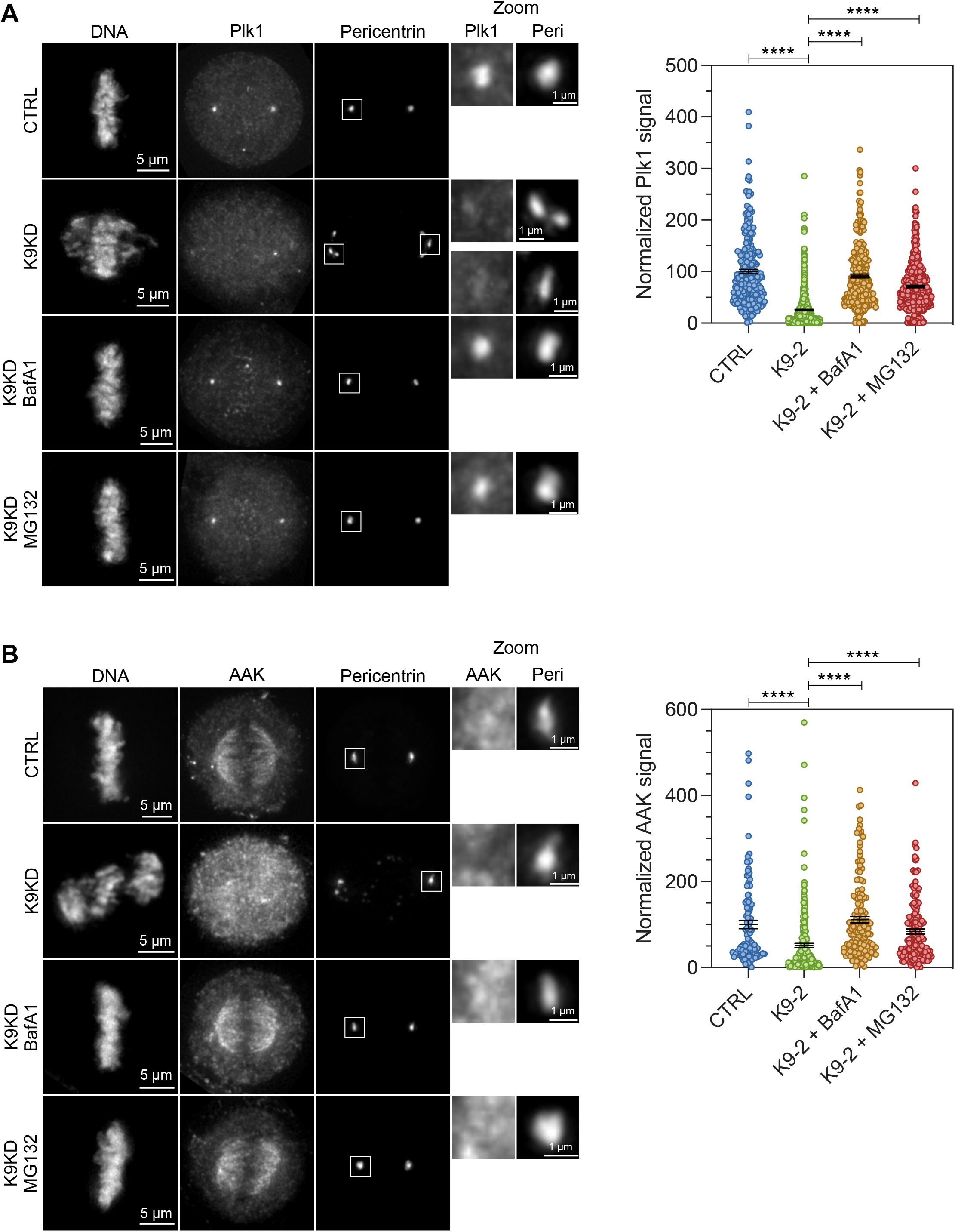
Kif9 depleted cells show lower levels of PLK1 and AURKA (AAK) at the centrosomes during mitosis. **(A)** HeLa cells were transfected with either control or Kif9 siRNAs, treated or not with Bafilomycin A1 or MG132, fixed and stained with DAPI for DNA, and antibodies against PLK1 and Pericentrin (for centrosomes). Kif9 depleted cells showed lower levels of PLK1 at the centrosome; levels that can be restored with either Bafilomycin A1 or MG132. Graph on the right shows the quantification of PLK1 levels at the centrosome. Data from at least 200 centrosomes per condition from 4 independent experiments and expressed as mean ± s.e.m. We used an unpaired t test as statistical analysis and the GraphPad Prism nomenclature where p values are categorized as follow 0.1234 (ns), 0.0332 (*), 0.0021 (**), 0.0002 (***), <0.0001 (****). Scale bar: 5 µm for the regular panels and 1 µm for the zoom panels. **(B)** HeLa cells were transfected with either control or Kif9 siRNAs, treated or not with Bafilomycin A1 or MG132, fixed and stained with DAPI for DNA, and antibodies against AURKA (AAK) and Pericentrin (for centrosomes). Kif9 depleted cells showed lower levels of AURKA at the centrosome; levels that can be restored with either Bafilomycin A1 or MG132. Graph on the right shows the quantification of AURKA levels at the centrosome. Data from at least 100 centrosomes per condition from 2 independent experiments and expressed as mean ± s.e.m. We used an unpaired t test as statistical analysis and the GraphPad Prism nomenclature where p values are categorized as follow 0.1234 (ns), 0.0332 (*), 0.0021 (**), 0.0002 (***), <0.0001 (****). Scale bar: 5 µm for the regular panels and 1 µm for the zoom panels.

**Figure 5.**
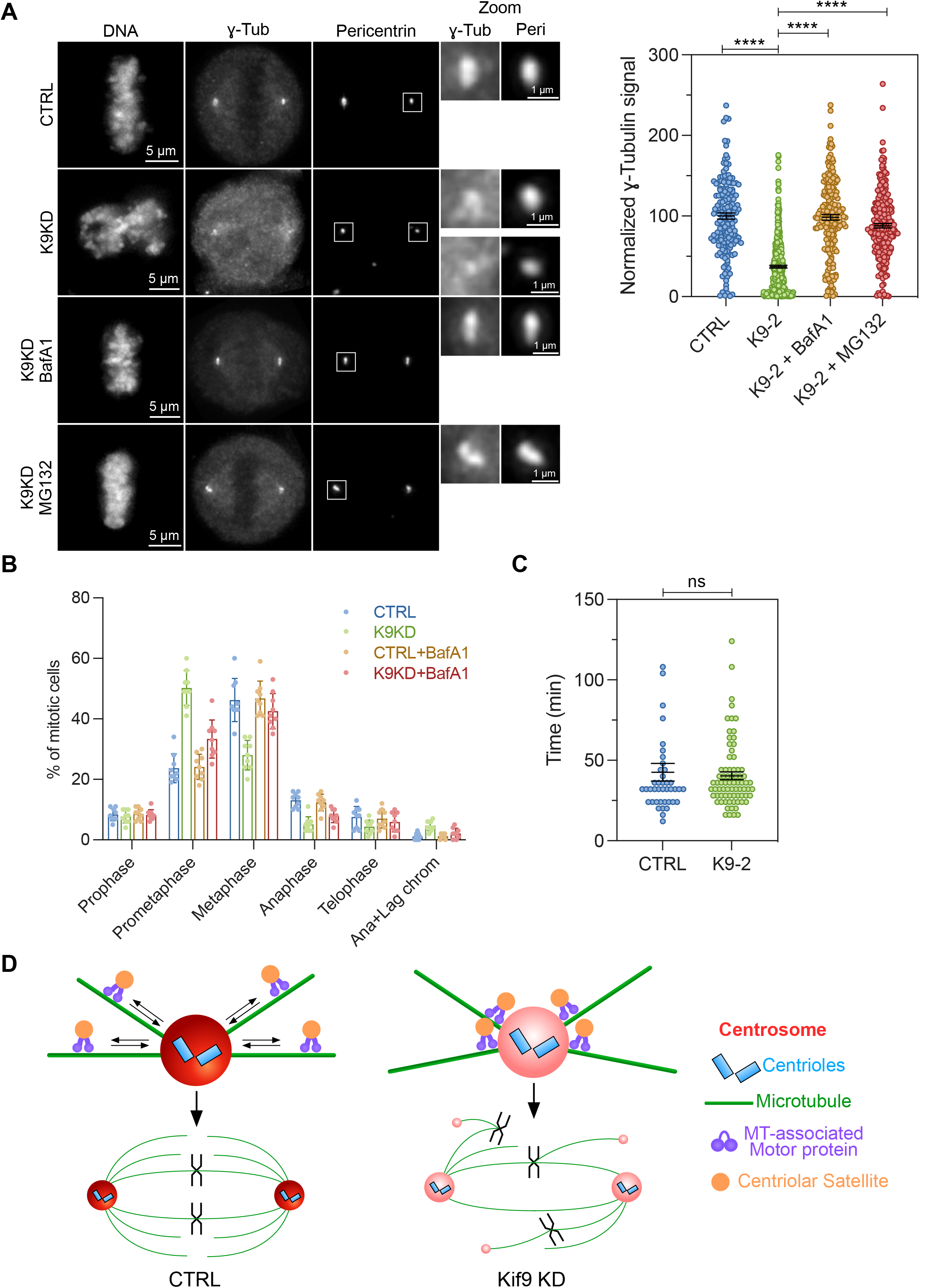
Kif9-depleted cells present lower levels of ɣ-tubulin at centrosomes in mitosis. Use of Bafilomycin A1 or MG132 recover the Kif9-KD phenotypes. **(A)** HeLa cells were transfected with either control or Kif9 siRNAs, treated or not with Bafilomycin A1 or MG132, fixed and stained with DAPI for DNA, and antibodies against ɣ-tubulin and Pericentrin (for centrosomes). Kif9 depleted cells showed lower levels of ɣ-tubulin at the centrosome; levels that can be restored with either Bafilomycin A1 or MG132. Graph on the right shows the quantification of ɣ-tubulin levels at the centrosome. Data from at least 150 centrosomes per condition from 3 independent experiments and expressed as mean ± s.e.m. We used an unpaired t test as statistical analysis and the GraphPad Prism nomenclature where p values are categorized as follow 0.1234 (ns), 0.0332 (*), 0.0021 (**), 0.0002 (***), <0.0001 (****). Scale bar: 5 µm for the regular panels and 1 µm for the zoom panels. **(B)** Quantification of the percentage of cells in every mitotic stage in fixed cells (spindle profile). HeLa cells were transfected with control siRNA, Kif9 siRNA, and treated or not with Bafilomycin A1. Cells were then fixed and stained with DAPI for DNA, and antibodies against tubulin (for MTs) and Pericentrin (for centrosomes). Kif9 depleted cells showed an increase in the number of prometaphase and a decrease of metaphases, phenotype recovered by the treatment with Bafilomycin A1. Cells lacking Kif9 also showed increased levels of anaphases with lagging chromosomes, also recovered by the treatment with the aforementioned drug. Data from n = 900 mitotic cells per condition from 3 independent experiments. Bars are mean ± s.e.m. **(C)** Mitotic timing from nuclear envelope breakdown (NEB) to the formation of a metaphase plate. Stable HeLa cells expressing H2B-GFP were transfected with either control or Kif9 siRNAs, treated with MG132 and recorded for 24 hours with 4 minutes steps. Kif9-depleted cells treated with MG132 had similar congression times as control cells. Data from at least 40 cells from 4 independent experiments. Bars are mean ± s.e.m. We used an unpaired t test as statistical analysis and the GraphPad Prism nomenclature where p values are categorized as follow 0.1234 (ns), 0.0332 (*), 0.0021 (**), 0.0002 (***), <0.0001 (****). **(D)** Model of how loss of Kif9 alter CS localization, centrosome maturation and triggers mitotic defects. During late G2/early mitosis CS in control cells move back and forth from the centrosomes to keep a balance between the amount of proteins acquired and the amount of proteins degraded at the centrosome. This balance allows regular PLK1 and AURKA activity that in turn results in proper centrosome maturation (dark red centrosome) that will provide this organelle with the structural integrity necessary to resist the action of spindle forces produced during chromosome movement in mitosis. In Kif9-depleted cells CS are closer to the centrosome providing this organelle with extra proteolysis-related factors that will degrade PLK1 and AURKA impeding proper centrosome maturation (light red centrosome). In this situation the spindle forces produced during mitosis fragment the PCM and create extra spindle poles that result in chromosome congression and segregation problems and increased mitotic timing.

### Kif9 depletion phenotypes in mitosis can be rescued by inhibiting protein degradation and autophagy

Several studies during the last decade have suggested that besides being the MTOC of the cell, the centrosome acts as a hub and scaffold for protein degradation through the proteasome and aggresome, and it has an essential function in autophagy ^2,3,50^. It has also been suggested that CS, at a distance from the centrosome, keep protein degradation machinery away from centrosome to prevent untimely ubiquitination and degradation of centrosomal components ^14^. In fact, CS have been linked to ubiquitination, autophagy and the control of proteolysis in the cell ^50–56^. Consequently, we believe that positioning CS closer to the centrosome in Kif9 depleted cells, in turn, positions the proteolytic machinery closer to centrosome-associated substrates leading to excess proteolysis. To test our hypothesis we performed Kif9 knockdowns followed by a treatment with either the proteasome inhibitor MG132 or the autophagy inhibitor Bafilomycin A1. These treatments were able to rescue the PCM fragmentation phenotype and to restore appropriate protein levels of PLK1, AURKA and ɣ-tubulin at the centrosome of mitotic cells (Fig. 4A-B, Fig. 5A and Supplementary Fig. 5A). Bafilomycin A1 treated Kif9-KD cells presented normal spindle profiles with close to normal proportions of prometaphase and metaphase stages (Fig. 5B and Supplementary Fig. 5B). This treatment also rescued chromosome missegregation during anaphase (Supplementary Fig. 5C). The proteasome inhibitor MG132 is extensively used in the mitotic field to arrest cells in metaphase. Accordingly, we find that MG132 treatment rescues chromosome congression (Supplementary Fig. 5D). We also analyzed the timing of congression from nuclear envelope breakdown (NEB) to the formation of a metaphase plate. MG132 treated Kif9-KD cells had similar timing to form a metaphase plate as control cells (Fig. 5C and Supplementary Fig. 5E). These results show that inhibiting protein degradation rescues the defective mitotic phenotypes seen in Kif9 depleted cells.

## DISCUSSION

The centrosome plays important roles in cell proliferation, differentiation and development (reviewed in ^1,57^). Centrosome maturation is essential for centrosomal function, and loss of spindle pole integrity has been linked to the formation of multipolar spindles, and chromosome congression and segregation problems during mitosis ^8,38,58–63^. To perform these different functions the composition of centrosomes changes through the cell cycle. Thus, cells must maintain a fine balance between the delivery of centrosomal proteins and the fraction of proteins degraded at this organelle. Our results support a hypothesis in which the cell accomplishes this spatially, using the kinesin motor Kif9 to regulate CS positioning, effectively sequestering them away from the centrosomes (Fig. 2B-C). This avoids disproportionate centrosomal protein degradation and enables proper centrosomal maturation (Fig. 4 and 5), necessary for the subsequent formation of a normal mitotic spindle to achieve error-free mitosis. As far as we know, this is the first report of a plus-end directed motor regulating CS positioning. Kif9 is expressed at low levels in our tested cells lines (Supplementary Fig. 1B-C and Supplementary Fig. 2A) and according to databases like OpenCell (https://opencell.czbiohub.org/). Thus, it has not been detected in the previous proteomic analysis of CS ^32,33^. However, the present work shows Kif9 is a *bona fide* plus-end directed motor responsible for CS positioning.

Protein degradation at the centrosome is necessary for general centrosomal function, embryonic development, neural morphology, for the immunological response and cell fate and cell cycle progression ^3^. Positionally governable CS have the potential to balance protein delivery with degradation to maintain centrosome homeostasis and function ^64^. In fact, several proteolysis (20 E3 ubiquitin-ligases and 4 deubiquitinating enzymes) and autophagy related enzymes have been found in the interactome of centriolar satellites ^33^. Defects in protein degradation at the centrosome have also been linked to changes in its morphology. For example, the lack of the E3-ubiquitin ligase HERC2 at the centrosome alters PCM morphology ^65^. Depletion of CS alters PCM composition promoting lower levels of Centrin, Pericentrin and Nek2 at centrosomes ^15,66^. It has also been previously reported that PCM1 sequesters MIB1 to CS to negatively control its activity ^56^. MIB1 and other proteasome related proteins also appear in our mass spectrometry data for Kif9 (Table 1). Interestingly, centrosomal levels of AURKA are elevated at the centrosome in PCM1 depleted cells ^67^, and cells exposed to the proteasome inhibitor MG132 for long periods of time exhibit increased protein levels at the centrosomes ^68^. These data show that proteins levels at the centrosome are highly regulated and that CS participate in this function. Matching these results, we show that positioning CS closer to the centrosome decreases levels of AURKA, PLK1 and ɣ-Tub, defects that can be rescued treating cells with MG132 or the autophagy inhibitor Bafilomycin A1 (Fig. 4A-B and Fig. 5A). All these data suggest that altering CS position by removing part of the motility machinery responsible for CS spatial positioning can affect the balance of recruitment/degradation of proteins at the centrosome, which is ultimately critical for centrosomal function (Fig. 5D).

## Acknowledgments

We thank Prof. Alex Merz at UW Biochemistry for his generous gift of that anti-GFP antibody. We also thank Dr. Paula Bucko and Dr. Kainat Khan for helpful discussions and creative feedback on the manuscript.

## Author contributions

Conceptualization: JJV, LW

Data Curation: JJV, LW

Investigation: JJV, MW

Data Analysis: JJV, JD, LW

Funding Acquisition: JJV, LW

Methodology: JJV, JD, MW, LW

Project Administration: LW

Software: JJV, JD

Supervision: JJV, LW

Validation: JJV, MW, LW

Visualization: JJV, LW

Writing – Original Draft Preparation: JJV, LW

Writing – Review & Editing: all authors;

Mass spectrometry experiments and analysis: AZ, MJM, TND

## Data and materials availability

All data are available in the main text or the supplementary materials. The mass spectrometry raw data is available at ProteomeXchange.

## MATERIALS AND METHODS

### Cell culture and plasmid transfection

HeLa, HCT116 and U2OS were purchased from ATCC and tested every month for mycoplasma contamination. No cell lines used for this study were found in the commonly misidentified cell lines database (ICLAC and NCBI Biosample). Cells were cultured in RPMI-1640 (Gibco/Thermofisher and Cytiva) with a 10% final concentration of FBS (Hyclone/Cytiva) in the presence of Pen-Strep (Corning). Media without antibiotics was used for siRNAs and plasmid transfections. Cells were maintained at 37 °C and 5% CO2. Cells were transfected with 2 µg of plasmid DNA, unless otherwise specified, using Nucleofector II (Lonza) according to manufacturer’s instructions.

### CRISPR cells lines

Kif9 CRISPR cell lines were made as indicated in Wagenbach et al^69^. The sequence for the guide RNA is the next: gRNA: 5’-ACCCATtctagctaaaaaga-3’.

### RNA interference

For Kif9 depletion, three different siRNAs targeted against three distinct regions of the Kif9-mRNA were used independently to assess Kif9-KD. siRNA#2: 5’-CCAACAGACAGACUGGUCGtt-3’ (sense), 5’-CGACCAGUCUGUCUGUUGGtt-3’ (antisense) (Ambion by Life Technologies); siRNA#3: 5’-GAAUUUCCAGAAGUCACUAtt-3’ (sense), 5’-UAGUGACUUCUGGAAAUUCag-3’ (antisense) (Ambion by Life Technologies); siRNA#4: 5’-AUAGCGUUUCUUCUAACUGGGCAGC-3’ (Invitrogen). siRNAs #2 and #4 significantly decreased the mRNA levels to the greatest extent and both had the same mitotic phenotypes. Thus, the siRNA#2 was used in all further experiments (unless otherwise indicated). Cells were transfected with 50 nM control (negative control siRNA#1 from Ambion) or Kif9 using Lipofectamine RNAiMax (Invitrogen) according to manufacturer instructions and incubated for 36-48 hours prior to experiments.

### Q-RT-PCR

Total cell RNA was isolated using a RNeasy mini kit (Qiagen). Retro-transcription was conducted using a Quantitect Reverse Transcription kit (Qiagen) with a first step of genomic DNA elimination. Q-PCR reactions were done using the SYBR green PCR kit (Qiagen) according to manufacturer’s instructions and an Applied Biosystems 7000 sequence detection system thermocycler. Primers for Kif9 detection: 5’-TGACCTATGACCCCATGGAT-3’ and 5’-CAGAAATGGCTGCAAAGTCA-3’. Samples lacking cDNA or RNA (no nucleic acids), and samples with just RNA were used as negative control, with no signal detected in both cases. As internal control we used GAPDH primers from Qiagen: Hs_GAPDH_1_SG QuantiTect Primer Assay (QT00079247).

### Drug treatments

For relocalization experiments, cells were transfected with either eGFP-FKBP-Kif9 (relocalizable form of Kif9) or eGFP-FKBP-Kif9 plus SH4-FRB-BFP (to allow relocalization to regions close to the cell membrane). Rapamycin at 1 µM final concentration was added 24h after transfection and incubated O/N at 37 °C. Cells were then fixed and processed for immunofluorescence. To recover the phenotypes of Kif9-KD, cells were treated with either 10 µM MG132 (Sigma) for 2h at 37 °C or 400 nM Bafilomycin A1 (APExBIO) for 4h at 37 °C after completion of siRNA treatment. Cells were then fixed and processed for immunofluorescence. For congression in MG132 experiments, cells were treated with siRNAs and afterward change to CO_2_ independent media (Gibco/Thermofisher) – 10% FBS supplemented with MG132 at 10 µM.

### Immunofluorescence

For fixed cells experiments, cells were plated in 18 mm diameter #1.5 glass coverslips (Electron Microscopy Sciences) and fixed in either 2% paraformaldehyde in methanol at −20 °C for 10 min or 4% paraformaldehyde in PBS buffer at 37 °C for 15 min depending on the antibody. Cells fixed with 4% PFA in PBS were then permeabilized with 0.5% Triton-X100, 5 minutes at room temperature. Coverslips were then blocked with 20% donkey serum for 1 h (Jackson Immunoresearch). Cells were then labeled with the next primary antibodies: mouse-anti-α-tubulin (1:500; Sigma monoclonal DM1α), rabbit-anti-PCM1 (1:250, ThermoFisher), mouse-anti-γ-tubulin (1:1,000; sigma GTU-88), rabbit-anti-pericentrin (1:500, Abcam), mouse-anti-pericentrin (1:500, abcam), mouse-anti-Plk1 (1: 200; Santa Cruz), mouse-anti-AURKA (1:250, Proteintech), mouse-anti-phospho-(T210)-Plk1 (1:200, abcam), rabbit-anti-phospho(T288)-AURKA (1:100, Cell Signaling), mouse-anti-Cep170 (1:250, Invitrogen), mouse-anti-CenpE (1:500, Santa Cruz). All the antibodies were applied overnight at 4 °C. Alexa-fluor 488, 568 and 647 secondary antibodies against mouse and rabbit IgG were used to detect the forementioned primary antibodies (Molecular Probes and Jackson Immunoresearch) The secondary antibodies were applied 2 hours at room temperature. Coverslips were mounted using ProLong® Diamond containing DAPI (Molecular Probes, Eugene, OR) and cured for 24 h. Cells used for super-resolution microscopy using SIM were mounted in ProLong® Diamond without DAPI.

### Microscopy

Fluorescent images were collected as either 0.25-μm or 0.5-μm Z-stacks on a Personal Deltavision microscope system (Applied Precision/GEHealthcare, Issaquah, WA) using a 60x 1.42 NA lens (Olympus, Tokyo, Japan). Images were deconvolved using SoftWorx 5.0 (Applied Precision/GEHealthcare). Alternatively, a GE InCell 2500 (GEHealthcare) system was used to capture images with the same conditions. Representative images are presented as a flat Maximum intensity Z-projection. Images projected as Sum Slices were used for quantification of fluorescence signal. Quantification of fluorescence intensity, co-localization and area occupied by CS were performed using the free programs CellProfiler and Fiji.

Super-resolution Structural Illumination Microscopy (SIM) imaging was performed on a Deltavision OMX V4 SIM microscope (GE Healthcare) equipped with an Olympus 60×/1.42, Plan Apo N objective, and a pco.edge sCMOS camera (chip size 2560 × 2160, and pixel size of 6.5 × 6.5 μm). Z-sections images were taken with 0.125 μm spacing. Images were reconstructed using softWoRx software from GE Healthcare.

For live-cell microscopy, HeLa cells stably expressing H2B-GFP were plated in Ibidi 35 mm HiQ4 4 chamber glass bottom imaging dishes to visualize nuclei during cell cycle. Immediately before imaging, regular RPMI media was replaced with CO_2_-independent-media – 10% FBS. Images were acquired on a Nikon Biostation IM-Q (Nikon) with a 20X lens as stacks of 4×3-µm Z-steps every 4 or 5 min for 24h. Alternatively, cells were cultured in Ibidi glass-bottom 24 multi-well plates. After changing media to CO_2_-independent media – 10% FBS, cells were imaged using a GE InCell 2500 system with a 20x lens. Time-lapse images were collected as 15 µm Z-stack series, with 3 µm Z-spacing, and with 4 or 5 min time intervals for 24 h. Images were deconvolved using the InCell software and processed with Fiji. Time-lapse movies were analyzed by hand using Fiji.

### Kif9 motility assays using TIRF microscopy

Kif9 fused to GFP (eGFP-Kif9) protein was expressed and purified from SF9 insect cells (Thermofisher). MTs purified from bovine brain tubulin were used for all motility experiments. To determine Kif9 motility direction we used dynamic MTs where 0.05 mg/mL GMPCPP Cy5-labelled MT seeds were attached to the coverslip using a rigor kinesin. Then, dynamic MTs were grown from the seeds by flowing into the chamber 1.0 mg/ml Alexa-Fluor-568 labelled tubulin. To calculate Kif9 motility speed we used live MTs without Cy5-seeds labelled with AlexaFluor-568. Sheared unlabeled dynamic MTs were bound to PEG-silane coated glass coverslip with rigor kinesin. Unbound MTs were displaced from flow chamber with one volume of 1 mg/mL Alexa568-labeled monomeric tubulin + 10 nM EGFP-Kif9 in 0.32x SQ buffer + 1 mM ATP + 1 mM GTP + antifade (1x SQ = 40 mM PIPES-K pH 6.85 = 100 mM K-acetate + 20 mM KCl + 4 mM MgCl2 + 1 mM EGTA + 0.063% Brij-35). EGFP-Kif9 motility was observed in TIRF on the growing red plus-end extensions of MTs using a Personal Deltavision microscope (Applied Precision) with 4-laser TIRF capabilities and an Olympus 60X 1.49 NA TIRF objective at 37 °C.

### Immunoprecipitation and western-blot

Immunoprecipitation for mass spectrometry. Protein A magnetic beads (New England Biolabs) were saturated with rabbit-anti-GFP, crosslinked with dimethyl pimelimidate, then quenched with ethanolamine. EGFP-Kif9 stably transfected cells and parental HCT116 cells grown with 0.1 µg/mL colcemid for 15 hours were washed off plates by trituration without trypsin (mitotic cells) then lysed, while all cells from similar plates grown without colcemid were lysed on ice in NP-40 buffer (0.25% NP-40 +150 mM NaCl + 25 mM HEPES-NaOH pH 7.5 + 11% glycerol + 0.5 mM EDTA + 2 mM MgCl2 + 1x HALT protease inhibitor (Pierce) + 1 mM NaPPO7 + 5 mM NaF + 1 mM betaglycerophosphate + 1 mM NaVO4 + 1 mM PMSF), homogenized by passing through a 25G neeedle, then centrifuged 10’ at 15,000g, 4 °C. Supernatant was incubated with anti-GFP beads 90’ at 4 °C, washed 1x with NP-40 buffer, then washed 2x with same buffer except NP-40 replaced by 1% PPS detergent (Expedeon). Immunoprecipitated complexes were released from beads with 0.2 M acetic acid and dried.

Immunoprecipitation for western-blot. HeLa cells were transfected with 2 µg of either a eGFP-Kif9 plasmid or the empty plasmid pEGFP-C1. After 40 hours, cells were collected and lysed. Anti-GFP beads and HeLa cell lysate in NP-40 buffer prepared as above. Washed beads were eluted by incubation in 1x NuPAGE sample buffer @ 37 °C, eluate was incubated at 70 °C then run on 4-12% NuPAGE gell and transferred to LF PVDF membrane (BioRad). Western-blot analysis were done with the next antibodies: rabbit-anti-PCM1 (1:250, ThermoFisher), mouse-anti-Cep170 (1:250, Invitrogen), mouse-anti-GAPDH (1:5,000, Calbiochem clone 6C5), and rabbit-anti-GAPDH (1:5,000, Sigma). Secondary antibodies fused to HRP and ECL (ThermoScientific) substrate were used to visualize PCM1 and Cep170. Secondary antibodies fused to AlexaFluor 568 were used to detect GAPDH: Goat-anti-Rabbit-Alexa568 (Molecular Probes/Invitrogen) and Donkey-anti-mouse-Alexa568 (Molecular Probes/Invitrogen).

### Mass spectrometry

Immunoprecipitated proteins were placed in a speedvac at room temperature for 4 hours until dry. Proteins were resuspended in 20 µL of 50 mM ammonium bicarbonate and pH was confirmed to be over 7. Samples were reduced in 10 mM final concentration DTT followed by incubation at 37°C for 30 minutes. Proteins were alkylated using 16 mM iodoacetamide at room temperature for 20 minutes in the dark followed by trypsin digestion with 0.4 µg Promega sequence grade modified trypsin for 6 hours at 37°C in an Eppendorf Thermomixer with shaking at 1000 rpm. Digestion was stopped by the addition of 250 mM HCl. Peptides were centrifuged at maximum speed in a bench-top microfuge for 10 mins and supernatant transferred to autosampler vials and stored at −80°C prior to analysis.

Digested peptides were analyzed by loading 3 µL onto a 150 µm Kasil fritted trap packed with 2 cm of ReprosilPur C18AQ (3 µm bead diameter, Dr. Maisch) at a flow rate of 2 µL per min. Reversed phase separation was done on a self-packed 75 µm i.d. 30 cm column and peptides were eluted at 0.25 µL/min using an acetonitrile gradient. A QExactive HF (Thermo Fisher Scientific) was used to perform MS in data dependent mode. Acquired mass spectra were converted into mzML format using ProteoWizard’s msConvert^70^. Peptides present in the samples were identified using Comet^71^ followed by post-processing using Percolator^72^. All data presented were filtered at a Percolator assigned q-value of ≤ 0.05. Comet search results were converted to Limelight XML and analyzed using the Limelight web application^73^.

Raw and processed mass spectrometry data are available via Limelight at: https://limelight.yeastrc.org/limelight/p/kif9-mitosis. In addition, complete search algorithm configuration files, fasta search databases, raw search output and raw MS data files were deposited to the ProteomeXchange Consortium via the PRIDE^74^ partner repository with the dataset identifier PXD048536.

### Statistical analysis and reproducibility

Statistical analysis was performed using Prism 9. We used Student’s t-test to compare the significance of the mean of two data sets. Experiments were repeated at least 3 independent times (unless otherwise indicated in the figure legends). We used the GraphPad Prism nomenclature where p values are categorized as follow 0.1234 (ns), 0.0332 (*), 0.0021 (**), 0.0002 (***), <0.0001 (****).

---

**Supplementary Figure 1.**
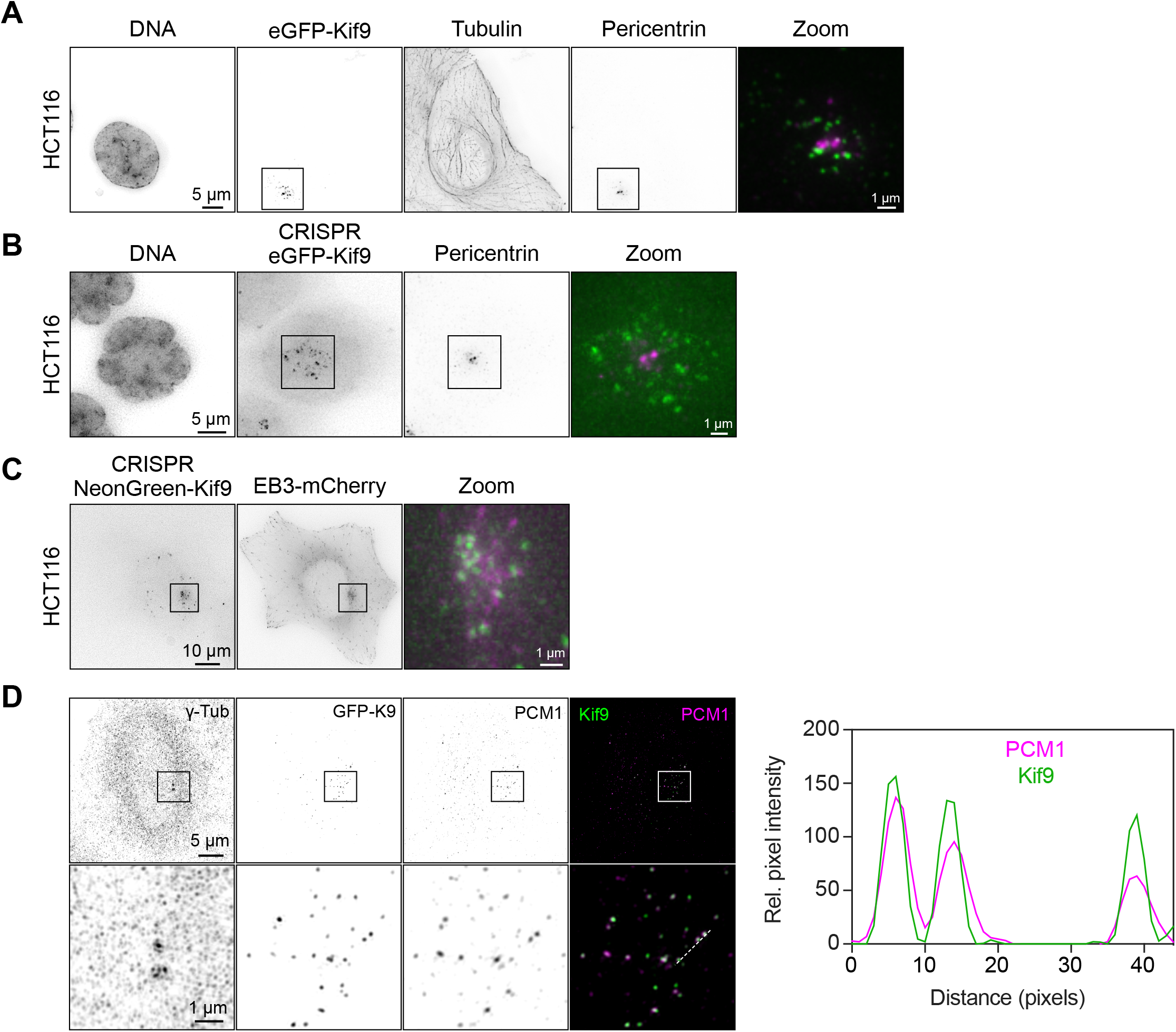
Kif9 localization in HCT116 cells and co-localization with PCM1 using SIM. **(A)** eGFP-Kif9 localizes to spot-like structures around the centrosome. HCT116 cells were transfected with a eGFP-Kif9 plasmid, fixed and stained with DAPI for DNA, and antibodies against Pericentrin (for centrosomes) and tubulin (for microtubules). Scale bar, 5 µm for the regular panels and 1 µm for the zoom panel. **(B)** eGFP-Kif9 localizes to spot-like structures around the centrosome. HCT116 cells were used to make CRISPR cell lines where eGFP-Kif9 is expressed from the endogenous Kif9 promoter. Cells were fixed and stained with DAPI for DNA, and antibodies against Pericentrin (for centrosomes). Scale bar, 5 µm for the regular panels and 1 µm for the zoom panel. **(C)** Neon-green-Kif9 localizes to spot-like structures around the centrosome. HCT116 cells were used to make CRISPR cells lines where Neon-green-Kif9 is expressed from the endogenous Kif9 promoter. Live cell images of the CRISPR cell line transfected with EB3-mCherry to label MTs tips and centrosome. Scale bar, 10 µm for the regular panels and 1 µm for the zoom panel. **(D)** Kif9-colocalizes with PCM1 using Structural Illumination Microscopy (SIM). HeLa cells were transfected with eGFP-Kif9, fixed and stained with antibodies against ɣ-tubulin (for centrosomes) and PCM1 (for CS). Graph Scale bar, 5 µm for the regular panels and 1 µm for the zoom panel. Graph on the right is a plot profile of the line in the magnified merge channel (bottom right panel).

**Supplementary Figure 2.**
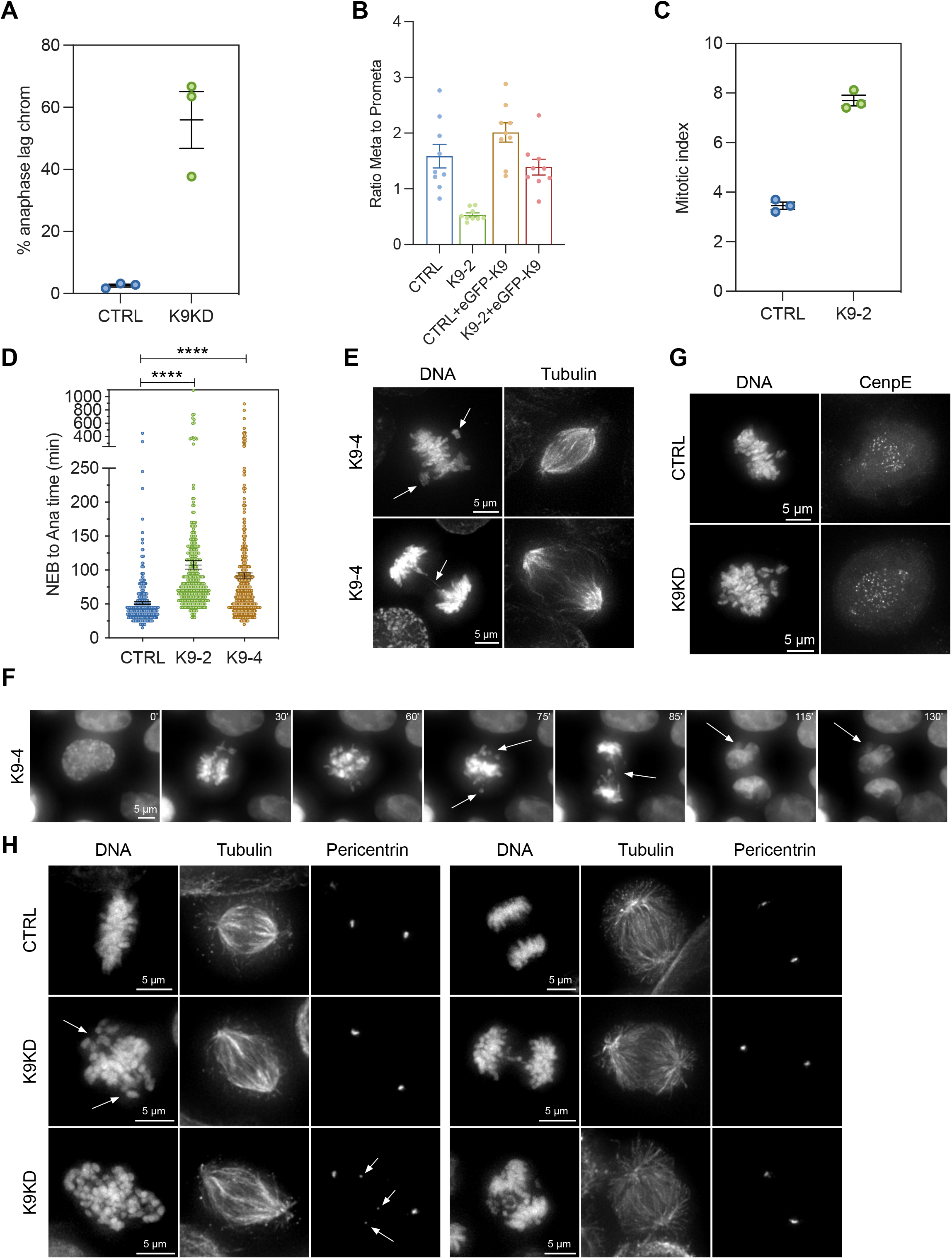
siRNAs against Kif9 reduce mRNA levels and Kif9-KD phenotype with siRNA#4. **(A)** Q-RT-PCR was performed as described in Methods to show Kif9 endogenous levels in HeLa cells and to show the efficiency of the knockdowns. This graph shows the Delta Rn (ΔRn), the magnitude of the signal generated by the given set of PCR conditions. In short, Rn is the fluorescent signal from emission of SYBR green divided by the emission of ROX dye (the passive reference dye). The ΔRn is the Rn value of an experimental reaction minus the Rn value of the baseline signal generated by the instrument. The dark lines correspond to the mean values of 3 experiments, and the light-colored area around the mean are the s.e.m. K9-2 = siRNA #2, K9-3 = siRNA#3 and K9-4 = siRNA#4. **(B)** Q-RT-PCR as shown in (a) but without reverse transcriptase (i.e. no cDNA in the reaction) as a negative control. No sample is amplified in this situation. The dark lines correspond to the mean values of 3 experiments, and the light-colored area around the mean are the s.e.m. K9-2 = siRNA #2, K9-3 = siRNA#3 and K9-4 = siRNA#4. **(C)** Ct values of the previous graphs showing a decrease in Kif9-mRNA levels (increased Ct values) in cells treated with Kif9 siRNA. K9-2 = siRNA #2, K9-3 = siRNA#3 and K9-4 = siRNA#4. **(D)** Relative mRNA expression of Kif9 mRNA from previous Q-RT-PCR graphs from 3 independent experiments. siRNAs #2 and #4 cause higher mRNA reduction than siRNA#3. **(E)** HCT116 cells treated with Kif9 siRNA#4 were fixed and stained with DAPI for DNA, and antibodies against PCM1 (for CS) and ɣ-tubulin (for centrosomes). Lack of Kif9 results in CS closer to the centrosome. Scale bar: 10 µm for the regular panels and 1 µm for the zoom panel.

**Supplementary Figure 3.**
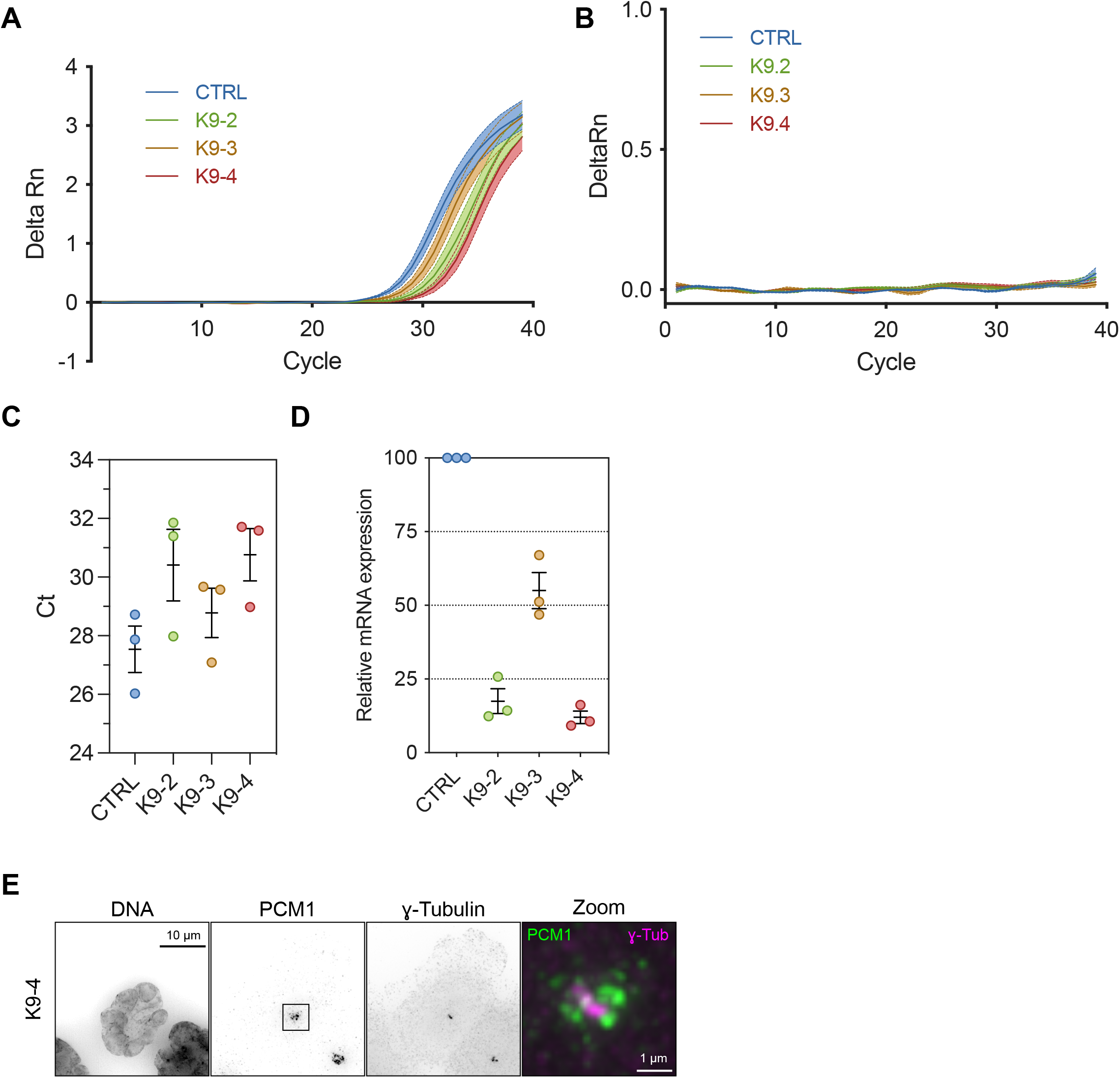
Cells depleted of Kif9 show a variety of mitotic phenotypes. **(A)** Quantification of lagging chromosomes in control and Kif9-depleted cells. Kif9-KD has higher number of lagging chromosomes. Data from 752 (control) and 314 (K9KD) anaphase HeLa cells from 3 independent experiments. **(B)** Quantification of the ratio metaphase to prometaphase cells from spindle profile in Fig. 3D. The decreased number of metaphase cells seen in the Kif9-KD can be recovered by expressing a siRNA resistant form of eGFP-Kif9. Data from 3 independent experiments is expressed as mean ± s.e.m. **(C)** Quantification of percentage of cells in mitosis respect to the total number of cells (i.e. mitotic index). HeLa cells depleted of Kif9 show higher number of cells in mitosis. Data from 3 independent experiments. **(D)** Quantification of mitotic timing from nuclear envelope breakdown (NEB) to anaphase onset (AO) from movies in Fig. 3E. Stable HeLa cells expressing H2B-GFP were transfected with either control or Kif9 siRNAs, and recorded for 24 hours with 5 minutes steps. Kif9 depleted cells show increased mitotic timing (CTRL = 51.30±2.23 min, K9-2 = 107.5±6.3 min, Kif9-4 = 91.35±4.49 min). Data from n = 312 control, 313 Kif9-2 and 441 Kif9-4 cells from 3 independent experiments. Data is shown as mean ± s.e.m. We used an unpaired t test as statistical analysis and the GraphPad Prism nomenclature where p values are categorized as follow 0.1234 (ns), 0.0332 (*), 0.0021 (**), 0.0002 (***), <0.0001 (****). **(E)** HeLa cells treated with Kif9 siRNA#4 were fixed and stained with DAPI for DNA, and antibodies against tubulin (for MTs). Lack of Kif9 results in chromosome congression and segregation problems similar to the ones seen with the siRNA#2. Scale bar: 5 µm. **(F)** Mitotic timing from nuclear envelope breakdown (NEB) to anaphase onset (AO). Stable HeLa cells expressing H2B-GFP were transfected with Kif9 siRNA#4, and recorded for 24 hours with 5 minutes steps. Representative images of the time lapse showing that Kif9 depleted cells have increased mitotic timing with chromosome congression and segregation problems, and the formation of small independent nuclei after nuclear formation (arrows in Kif9-4 panels). Scale bar, 5 µm. **(G)** HeLa cells treated with Kif9 siRNA#2 were fixed and stained with DAPI for DNA, and antibodies against CenpE. Lack of Kif9 does not affect CenpE localization to kinetochores. Scale bar, 5 µm. **(H)** HCT116 cells were transfected with either control or Kif9 siRNAs, fixed and stained with DAPI for DNA, and antibodies against tubulin (for MTs) and Pericentrin (for centrosomes). Kif9 depleted cells showed congression problems (arrows in DNA channel) and fragmentation of the PCM (arrows in Pericentrin channel). They also present defects during chromosome segregation in anaphase (right panels). Scale bar, 5 µm.

**Supplementary Figure 4.**
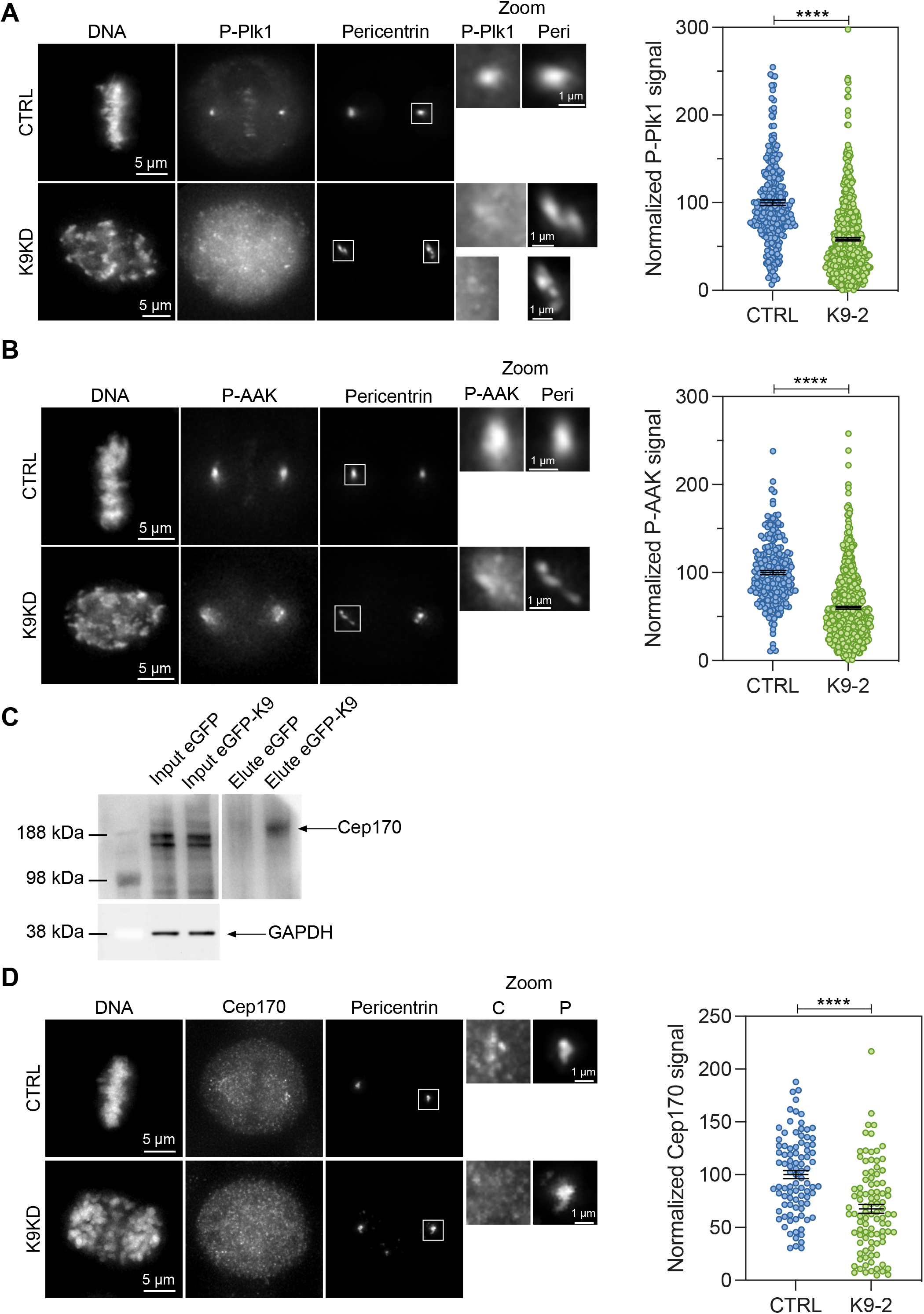
Levels of P-PLK1 and P-AURKA at the centrosome in Kif9-depleted mitotic cells and Cep170 interaction. **(a)** HeLa cells were transfected with either control or Kif9 siRNAs, fixed and stained with DAPI for DNA, and antibodies against P-PLK1 (T210) and Pericentrin (for centrosomes). Kif9 depleted cells showed lower levels of P-PLK1 at the centrosome. Graph on the right shows the quantification of P-PLK1 levels at the centrosome. Data from at least 200 centrosomes per condition from 3 independent experiments and expressed as mean ± s.e.m. We used an unpaired t test as statistical analysis and the GraphPad Prism nomenclature where p values are categorized as follow 0.1234 (ns), 0.0332 (*), 0.0021 (**), 0.0002 (***), <0.0001 (****). Scale bar: 5 µm for the regular panels and 1 µm for the zoom panels. **(B)** HeLa cells were transfected with either control or Kif9 siRNAs, fixed and stained with DAPI for DNA, and antibodies against P-AURKA (T288) and Pericentrin (for centrosomes). Kif9 depleted cells showed lower levels of P-AURKA at the centrosome. Graph on the right shows the quantification of P-AURKA levels at the centrosome. Data from at least 200 centrosomes per condition from 3 independent experiments and expressed as mean ± s.e.m. We used an unpaired t test as statistical analysis and the GraphPad Prism nomenclature where p values are categorized as follow 0.1234 (ns), 0.0332 (*), 0.0021 (**), 0.0002 (***), <0.0001 (****). Scale bar: 5 µm for the regular panels and 1 µm for the zoom panels. **(C)** Co-immunoprecipitation Kif9-Cep170. Immunoprecipitation of eGFP-Kif9 shows an interaction with Cep170. HeLa cells were transfected either with eGFP-Kif9 or the eGFP empty plasmid. Immunoprecipitation was performed with an anti-GFP antibody followed by immunoblotting with an anti-Cep170 antibody. **(D)** HeLa cells were transfected with either control or Kif9 siRNAs, fixed and stained with DAPI for DNA, and antibodies against Cep170 and Pericentrin (for centrosomes). Kif9 depleted cells showed lower levels of Cep170 at the centrosome. Graph on the right shows the quantification of Cep170 levels at the centrosome. Data from at least 90 centrosomes per condition from 3 independent experiments and expressed as mean ± s.e.m. We used an unpaired t test as statistical analysis and the GraphPad Prism nomenclature where p values are categorized as follow 0.1234 (ns), 0.0332 (*), 0.0021 (**), 0.0002 (***), <0.0001 (****). Scale bar: 5 µm for the regular panels and 1 µm for the zoom panels.

**Supplementary Figure 5.**
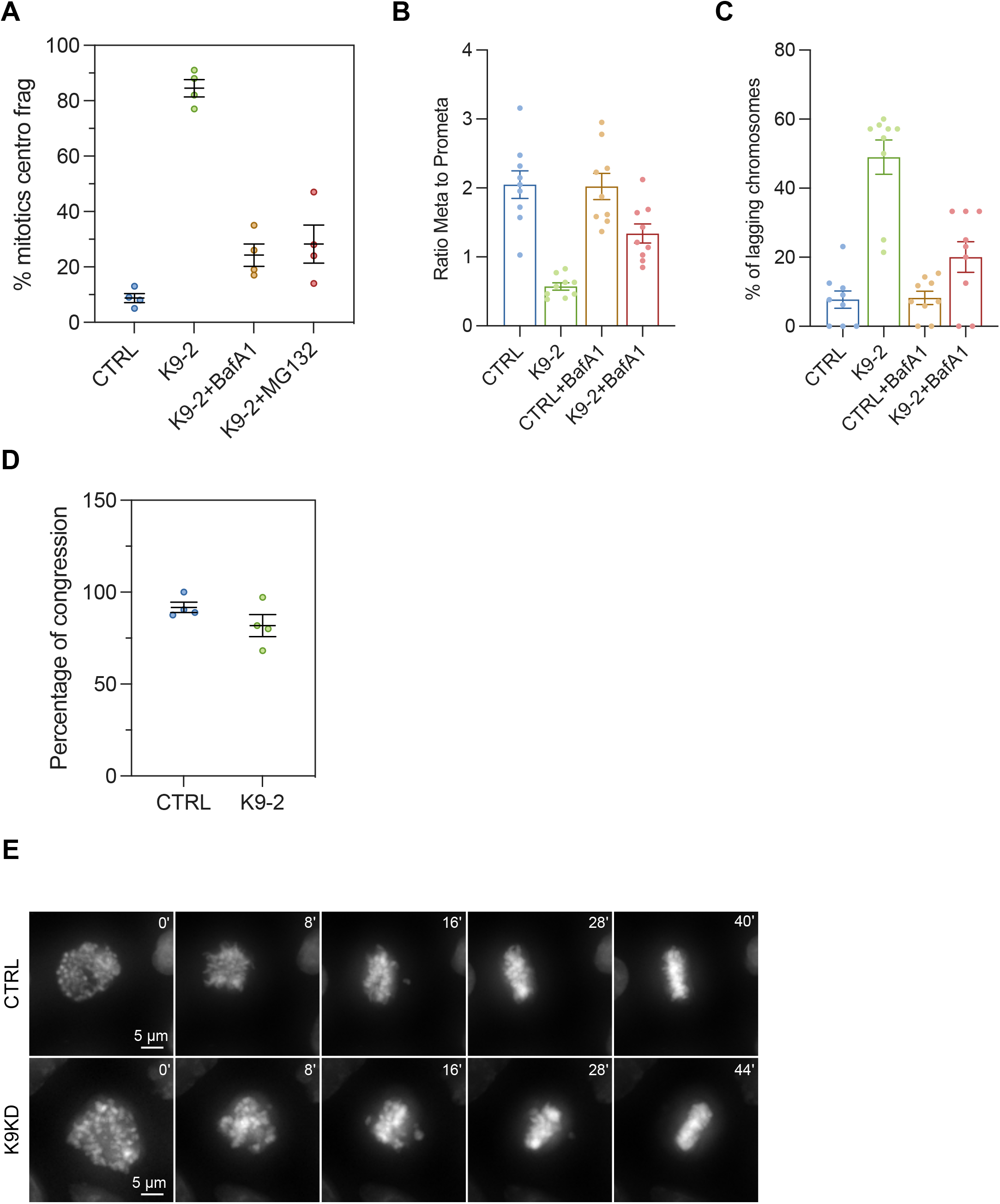
Recovery of Kif9-KD mitotic phenotypes using Bafilomycin A1 or MG132. **(A)** Quantification of PCM fragmentation in HeLa mitotic cells transfected with either control or Kif9 siRNA, treated or not with Bafilomycin A1 or MG132, fixed and stained with DAPI for DNA, and antibodies against PLK1 and Pericentrin (for centrosomes). Kif9 depleted cells showed increased PCM fragmentation that can be restored with either Bafilomycin A1 or MG132. Data from 400 mitotic cells per condition from 4 independent experiments and expressed as mean ± s.e.m. **(B)** Quantification of the ratio metaphase to prometaphase cells from spindle profile in Fig. 5B. The decreased number of metaphase cells seen in the Kif9-KD can be recovered by Bafilomycin A1. Data from 3 independent experiments is expressed as mean ± s.e.m. **(C)** Quantification of the percentage of lagging chromosomes in anaphase cells from spindle profile in Fig. 5B. The increased number of lagging chromosomes seen in the Kif9-KD can be recovered by Bafilomycin A1. Data from 3 independent experiments is expressed as mean ± s.e.m. **(D)** Percentage of cells that congress and form a metaphase plate in HeLa-H2B-GFP cells treated with either control or Kif9 siRNA and MG132 and recorded for 24 hours with 4 minutes steps. Kif9-depleted cells treated with MG132 had similar percentage of congression as control cells. Data from at least 40 cells from 4 independent experiments. Bars are mean ± s.e.m. **(E)** Mitotic timing from nuclear envelope breakdown (NEB) to the formation of a metaphase plate. Stable HeLa cells expressing H2B-GFP were transfected with either control or Kif9 siRNAs and treated with MG132. Cells were recorded for 24 hours with 4 minutes steps. Representative images of the time lapse showing that Kif9 depleted cells have similar mitotic timing as control cells in the presence of MG132. Scale bar, 5 µm.

